# Relaxed DNA substrate specificity of transposases involved in programmed genome rearrangement

**DOI:** 10.1101/2025.03.17.643836

**Authors:** Matt W.G. Walker, Takahiko Akematsu, Erhan Aslan, Danylo J. Villano, Harrison S. Fried, Hui Lan, Samuel H. Sternberg, Laura F. Landweber

## Abstract

During post-zygotic development, the ciliate *Oxytricha trifallax* undergoes massive programmed genome rearrangement that involves over 225,000 DNA cleavage and joining events. An *Oxytricha* family of Tc*1/mariner* transposons, known as Telomere-Bearing Elements (TBEs), encodes a transposase that has been implicated in rearrangement, but its high copy number (>34,000 paralogs) has precluded genetic strategies to investigate its DNA recognition properties directly in *Oxytricha*. Here, we developed a heterologous strategy to assay TBE transposase expression and activity in *E. coli*, revealing highly promiscuous DNA cleavage properties. Systematic ChIP-seq experiments allowed us to define the DNA binding specificities of multiple distinct transposase subfamilies, which exhibited a binding and cleavage preference for short, degenerate sequence motifs that resemble features present within the TBE transposon ends. The relaxed sequence preference is striking for autonomous transposases, which typically recognize their end sequences with strict specificity to avoid compromising host fitness. Finally, we developed a custom antibody to investigate TBE transposases in their native environment and found that they precisely localize to the developing nucleus exclusively during the rearrangement process. Collectively, this work establishes a robust heterologous workflow for the biochemical investigation of enzymes that have been repurposed for large-scale genome rearrangements.

## INTRODUCTION

The genomes of many organisms undergo programmed rearrangements that require precise DNA elimination and assembly (Smith *et al*., 2020; Wang & Davis, 2014). Arguably the most complex programmed genome rearrangements occur in the ciliated protozoan, *Oxytricha trifallax*. Like most ciliates, *Oxytricha* is binucleated, with a separate germline micronucleus (MIC) and somatic macronucleus (MAC) (Prescott, 1994). During the vegetative (asexual) life cycle, the MIC genome replicates through mitosis, while the MAC genome divides through amitosis, with chromosomes imperfectly segregated to the daughter nuclei. During sexual reproduction, conjugating cells exchange haploid micronuclei that undergo syngamy, the parental MAC degrades and one copy of the zygotic germline MIC develops into a new MAC, initiating an intricate cascade of recombination events requiring over 225,000 DNA cleavage and joining events (X. Chen *et al*., 2014; Prescott, 1994; Swart *et al*., 2013). This rearrangement process also results in the full elimination of >90% of germline DNA.

The MIC is present in multiple copies per cell and is largely transcriptionally silent, in part because a large portion of its precursor gene segments are in the wrong order relative to the transcribed version in the MAC; individual gene segments are also interrupted by internally eliminated sequences (IESs) that must be removed in order for retained macronuclear-destined sequences (MDSs) to arrange to produce functional MAC genes (**Figure 1A**). In the MIC, MDSs are flanked by short direct repeats, called *pointers*, only one of which remains after rearrangement. MDS joining produces >16,000 new ‘nanochromosomes’ that each acquire two telomeres and are amplified to high copy number (average ploidy ∼1900), exhibit an average length of 3.2 kb, and usually encode only one gene (X. Chen & Landweber, 2016; Lindblad *et al*., 2019; Prescott, 1994; Swart *et al*., 2013). Previous studies identified key molecular components that are involved in the rearrangement process: 27-nt Piwi-interacting small RNAs (piRNAs) mark MDSs for retention (Fang *et al*., 2012), long noncoding RNAs transcribed from parental MAC chromosomes program the MDS assembly process (Nowacki *et al*., 2008), and a large family of germline-encoded transposases are essential for DNA processing (Nowacki *et al*., 2009).

**Figure 1.**
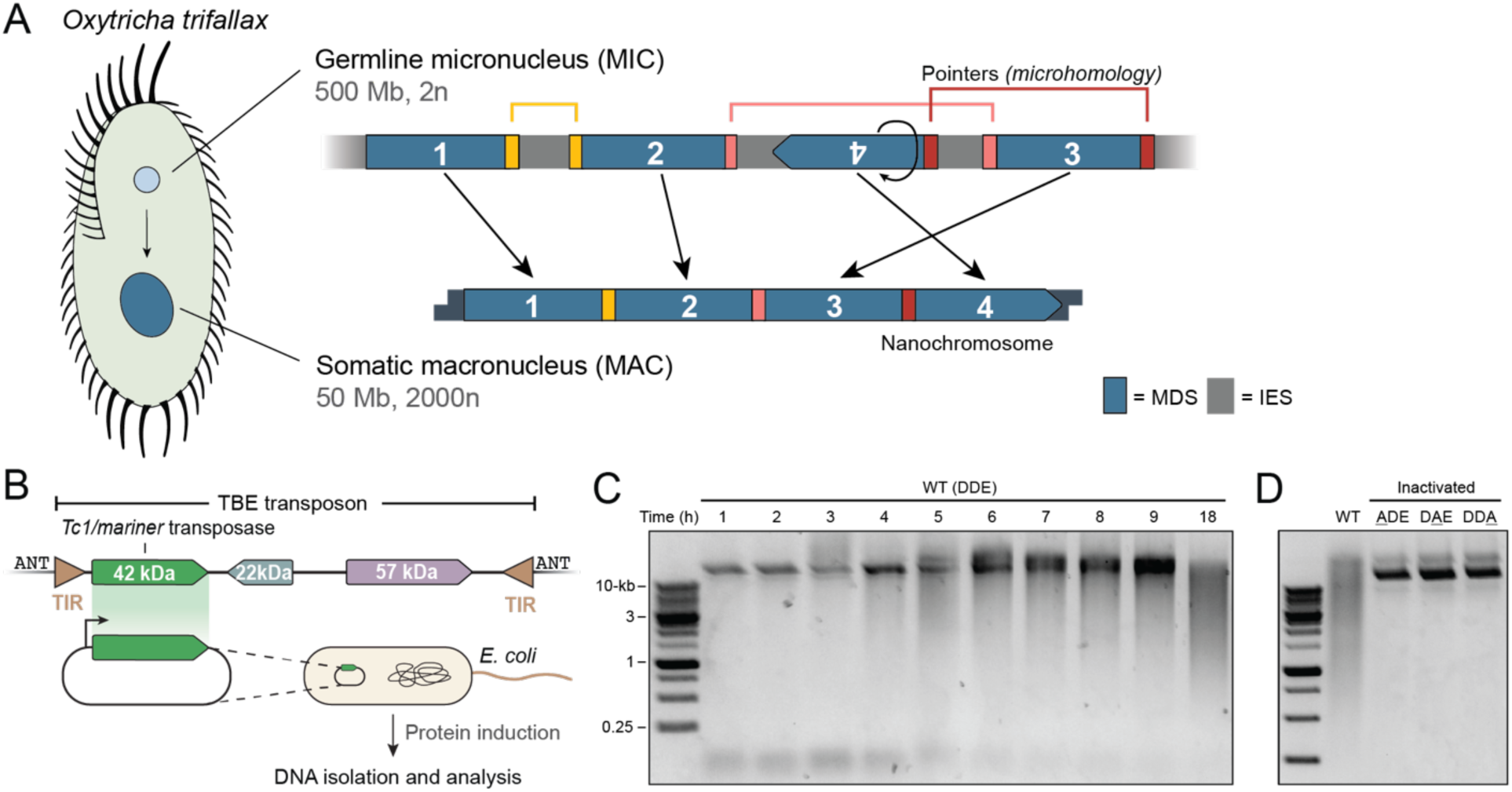
The TBE transposase induces DNA fragmentation in *E. coli*. (**A**) Genome rearrangement in *Oxytricha trifallax*. *Oxytricha* cells contain a germline micronucleus (MIC), which is diploid (2n) and encodes ∼500 Mb of genome content. The somatic macronucleus (MAC) is polyploid (2000n) and encodes ∼50 Mb of genome content. Blue rectangles illustrate macronuclear destined sequences (MDSs), which are separated by internally eliminated sequences (IESs, gray) in the MIC. During rearrangement, IESs are eliminated and MDSs rearrange at short homologous sequences (pointers, colored bars) to form functional gene-length nanochromosomes. (**B**) Schematic of TBE transposon genetic architecture (top), and heterologous TBE transposase expression approach in *E. coli*. Terminal inverted repeats (TIRs) are indicated by brown inverted triangles, which are flanked by the ANT target-site duplication. (**C-D**) DNA fragmentation analysis upon TBE transposase induction in *E. coli*, using either the WT transposase with intact DDE motif (**C**), or catalytically inactivated transposase mutants bearing either an <u>A</u>DE mutation or D<u>A</u>E or DD<u>A</u> (**D**). 500 ng DNA was separated by electrophoresis on a 1% agarose gel and visualized using SYBR Safe.

TBE, or Telomere-Bearing Element, transposons are abundant in the MIC, present at >34,000 copies that comprise almost 15% of the germline genome, but they are completely eliminated during development. TBEs encode three distinct ORFs, one of which is a 42 kDa transposase (**Figure 1B**). Phylogenetic analyses indicated that the TBE transposase belongs to the Tc*1*/*mariner* family (Doak *et al*., 1994), which uses a DDE catalytic triad to perform the DNA cleavage and strand joining chemistry required for transposition (Hickman & Dyda, 2016). Phylogenetic analyses also revealed that there are three superfamilies for the TBEs – TBE1, TBE2, and TBE3 – with TBE2 further subdivided into TBE2.1 and TBE2.2 (X. Chen & Landweber, 2016). The TBE moniker originates from the terminal inverted repeat (TIR) sequences flanking the elements, which encode telomere-like repeats (G_4_T_4_)_4_. Tc*1*/*mariner* family elements typically contain inverted repeats at their end sequences, which delineate the boundaries of the mobile element and recruit the transposition machinery with strict specificity during mobilization. In *Oxytricha*, TBEs are detected as extrachromosomal circular DNA (eccDNA) during genome rearrangement, indicating that they are excised as circular products (Williams *et al*., 1993; Yerlici *et al*., 2019), although direct mobilization activity has not been experimentally observed. While TBE transposons likely spread and proliferate via transposition, extant TBE transposases may instead play a predominant role in DNA excision, akin to the role of RAG1 recombinase in V(D)J recombination, more often removing rather than mobilizing genomic elements. Unlike Tc*1*/*mariner* elements, which recognize a TA target site and are flanked by two copies of that motif after integration, TBEs are flanked by an ANT motif, which is strikingly consistent with the most common motif of 3-bp pointer sequences (X. Chen *et al*., 2014). This overlap suggests that the motif is either a requirement for transposase-mediated DNA elimination, or a remnant of transposon integration events that led to the emergence of IESs (Klobutcher & Herrick, 1997).

Previous work on TBEs indicated their involvement in programmed DNA elimination. Specifically, RNAi knockdown of transposase gene families led to stalled rearrangement of MAC genomic loci, arrested development, and the accumulation of high molecular weight DNA along with incompletely or improperly rearranged genes (Nowacki *et al*., 2009). However, failed rearrangement only occurred when RNAi was directed against all 3 superfamilies of TBEs, limiting the utility of this approach to investigate the activity of specific TBE-encoded transposases. Furthermore, while both reverse and forward genetic tools in *Oxytricha* can target the MAC (Clay *et al*., 2019; Fang *et al*., 2012), techniques for genetic manipulation of the germline MIC are severely compromised by its low copy number and small size compared to the MAC, and this severely limits our ability to determine how and where TBE transposases cleave DNA to effect precise DNA elimination during development.

To circumvent these limitations, we developed a heterologous expression approach in *Escherichia coli*, which allowed us to gain insights into DNA binding and cleavage activity of the TBE transposase. We uncovered DNA binding preferences within the *E. coli* genome and its own TBE transposon ends, and find that this binding activity is consistent with DNA cleavage, which we detected using deep-sequencing to profile fragmented DNA. Our work presents the first direct evidence of TBE transposase binding and cleavage activity, and establishes a robust workflow for studying a previously intractable enzyme implicated in genome rearrangement.

## MATERIALS AND METHODS

### Plasmid and *E. coli* strain construction

Plasmids and oligonucleotides used in this study are listed in **Supplementary Tables 1** and **2**, respectively. *E. coli* codon-optimized open reading frames for TBE transposon-encoded proteins were synthesized and cloned into expression vectors under the control of an IPTG-inducible T7 promoter on medium-copy pET vectors with carbenicillin resistance. Epitope-tagged protein expression vectors were constructed by appending a C-terminal FZZ tag (3×FLAG-tag, TEV protease cleavage size, and ZZ domain of protein A, described in (Kataoka *et al*., 2010)). Catalytically-inactivated derivatives of the TBE transposase proteins were constructed by ‘around-the-horn’ PCR, to mutate each residue of the catalytic triad (DDE) to alanine (<u>A</u>DE, D<u>A</u>E and DD<u>A</u>). All plasmids were cloned into NEB Turbo cells (NEB), purified (QIAprep Spin Miniprep Kit), and verified by Sanger sequencing (GENEWIZ).

Four custom *E. coli* BL21(DE3) strains were generated encoding a genomically-integrated TBE transposon belonging to each of the TBE families (TBE1 from contig ctg7180000067530, TBE2.1 from contig 7180000067905, TBE2.2 from contig ctg7180000069065, and TBE3 from ctg7180000089059). These strains were generated by Himar1C9-mediated transposition. We cloned representative TBE transposons from the four TBE families onto a temperature-sensitive pSC101 plasmid, downstream of a kanamycin resistance cassette, which was flanked by *mariner*-family transposon end sequences. The plasmid also encoded a transposase expression cassette, which produced the Himar1C9 protein required for transposition of the *mariner* element. The temperature-sensitive plasmid was cured before selecting for antibiotic resistance, which could only be conferred after genomic integration of the TBE element. Specifically, after transformation by heat shock, cells were recovered at the restrictive temperature and plated on LB agar plates supplemented with kanamycin (50 µg/ml), and colonies that grew were confirmed by PCR to have lost the temperature-sensitive TBE plasmid. The genomic integration was later confirmed for each of the four strains by Illumina deep sequencing (Smear-Seq and ChIP-seq), which detected a single copy of the transposon in the genome and revealed the integration site, allowing us to generate complete *E. coli* reference genomes for each strain that were used for read mapping and analysis.

### DNA fragmentation assay and time course experiments

The DNA fragmentation analysis relied on inducing transposase expression in liquid media, performing a miniprep to isolate DNA, and separating DNA by electrophoresis on a 1% agarose gel. First, chemically competent *E. coli* BL21(DE3) cells were transformed and single colonies were grown overnight at 37 °C in LB medium with 100 µg/ml carbenicillin. The next day, cultures were diluted 1:100 in fresh LB medium with 100 µg/ml carbenicillin and grown at 37 °C until mid-logarithmic phase (OD_600_=0.4). Cultures were then induced with 0.1 mM IPTG and transferred to 16 °C. For the time course experiments, multiple 2 ml cultures were set up in replicate. After the defined induction time, cell density was measured (OD_600_), DNA was isolated (QIAGEN Plasmid Mini Kit), and DNA concentration was measured (DeNovix Spectrophotometer). 500 ng of DNA was loaded onto each lane of a 1% agarose gel stained with SYBR Safe (Thermo Fisher Scientific), subjected to electrophoresis (130 V for 20 min), and imaged (Bio-Rad GelDoc XR Imaging System).

### Structural predictions and sequence alignments

Structural models of the TBE transposase dimer were predicted using AlphaFold version 2.3.1 (commit 18e12d6) (Jumper *et al*., 2021), using the multimer model preset (--model_preset=multimer) (Evans *et al*., 2022), and displayed in ChimeraX (v1.6). Structural models of the TBE transposase dimer interacting with 40-bp of transposon end DNA were predicted using RoseTTAFoldNA (Baek *et al*., 2023). MSAs were generated by Clustal Omega run on EMBL’s Job Dispatcher (Madeira *et al*., 2022), and visualized in Jalview version 2.11.3.2 (Waterhouse *et al*., 2009).

### NGS approach for sequencing fragmented DNA (“Smear-seq”)

Our approach to sequencing fragmented DNA relied on DNA isolation, dA-tailing, second-strand synthesis, sonication, library prep, and next-generation sequencing (**Figure 2A**). First, the wildtype or catalytically-inactivated transposase was expressed as in the DNA fragmentation assay, wherein we diluted overnight cultures of *E. coli* BL21(DE3) cells expressing the transposase 1:100 in LB with 100 µg/ml carbenicillin, induced mid-logarithmic phase cultures with 0.1 mM IPTG, and grew cultures for 18 h at 16 °C. DNA was isolated (QIAGEN Plasmid Mini Kit), and A-tailing was performed using an approximate molar ratio of 1:1,000 of DNA:dATPs (NEB Terminal Transferase). After incubating at 37 °C for 30 min and heat-inactivating at 70 °C, DNA was cleaned and eluted in 40 µl Milli-Q H_2_O (QIAquick PCR Purification Kit). Concentration was measured by fluorometry (DeNovix dsDNA High Sensitivity Assay) before second-strand synthesis using oligo(dT)18 primers and PrimeSTAR Max DNA Polymerase (Takara). DNA was cleaned and eluted in 40 µl MilliQ H_2_O and the concentration was measured by fluorometry before normalized samples were loaded into a microTUBE AFA Fiber Crimp-Cap and TE buffer was added to a total volume of 130 µl. Samples were then sheared by sonication on an M220 Focused-ultrasonicator (Covaris) using the following SonoLab 7.2 settings: peak power, 50.0; duty factor, 20; cycles/burst, 200; treatment time, 150 s. After sonication, DNA was prepared for NGS using the NEBNext Ultra II DNA Library Prep for Illumina (NEB). After adaptor ligation, DNA was purified without size selection using AMPure XP beads (Beckman Coulter) and appended sequencing indexes with 10 cycles of PCR and NEBNext Q5 Master Mix. Two-sided size selection was used to clean and select ∼450 bp fragments. To first remove small fragments, 0.55× AMPure XP bead solution was added and samples were allowed to separate on a magnetic rack before the supernatant containing small DNA fragments was removed. Then, to remove large fragments, 0.35× AMPure XP bead solution was added to each sample, separated on a magnetic rack, and the supernatant discarded. Beads were then washed twice in 80% (v/v) EtOH and eluted in 12 µl of TE buffer. Concentration was determined by fluorometry (DeNovix dsDNA High Sensitivity Kit) and samples were sequenced using a NextSeq High Output Kit with 150 cycles (Illumina).

**Figure 2.**
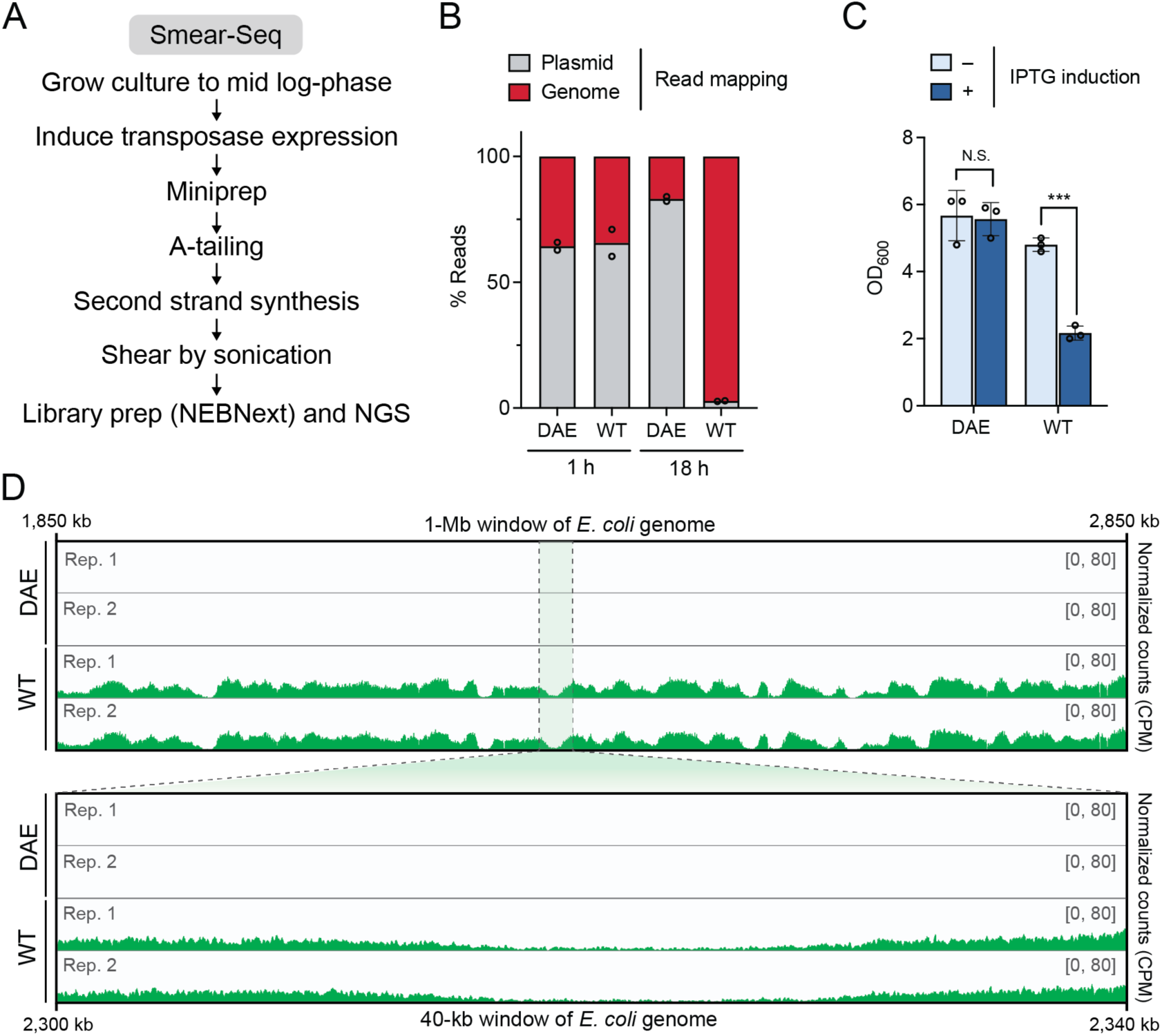
The TBE transposase cleaves genomic DNA and induces cell death. (**A**) ‘Smear-seq’ workflow for sequencing fragmented DNA in *E. coli*. (**B**) Percentage of reads from each sample mapping to the transposase expression plasmid or to the *E. coli* genome. Each point represents one independent biological replicate. (**C**) Optical cell density (OD_600_) of the WT or inactivated (DAE) transposase after 18 hours with and without IPTG induction. Data are shown as mean +/− SD for n=3 biological replicates; *** *p* < 0.001 (unpaired *t*-test); N.S., not significant. (**D**) Read coverage across the *E. coli* genome in a 1-Mb window (top) or 40-kb window (bottom). The y-axis represents read counts per million (CPM), with the axis limit set to 80 for each sample.

After sequencing, paired-end reads were filtered and trimmed using fastp (S. Chen *et al*., 2018) and mapped to the *E. coli* BL21(DE3) reference genome and to the TBE expression plasmid using bowtie2 (Langmead & Salzberg, 2012). Then, we used Samtools to sort, index, and filter multi-mapping reads (Danecek *et al*., 2021). Finally, coverage was normalized to counts per million (CPM) using the deepTools2 command bamCoverage (Ramírez *et al*., 2016).

### *Oxytricha* culture conditions and mating

*Oxytricha* strains were cultured and mated as in Fang *et al*., 2012. Two wild type *Oxytricha* strains with different mating types, JRB310 and JRB510 were cultured in Pringsheim media (0.11 mM Na_2_HPO_4_, 0.08 mM MgSO_4_, 0.85 mM, Ca(NO_3_)_2_, 0.35 mM KCl, pH 7.0) supplemented with *Chlamydomonas reinhardtii* and *Klebsiella* as the food source. To induce mating, cells were allowed to deplete *Chlamydomonas* overnight, filtered through cheesecloth to remove any remaining debris, and equal amounts of cells were mixed at a density of 5,000 cells/ml with a final volume of 300 ml. Pairs typically form 3–4 hours post-mixing, and maximum pairing efficiency (∼70–80%) is generally achieved 12 hours post-mixing.

### Production of anti-TBE transposase antibodies

The peptide KRQHLNSKPKRPLK was synthesized and used for immunization in rabbits to yield anti-TBE transposase antibodies (GenScript). Rabbits were immunized three times at two-week intervals. 4.28 mg antibody was purified and suspended in 7.5 ml PBS with 0.02% ProClin 300 preservative. The antibody was validated by Western hybridization in *E. coli* cells expressing the TBE transposase, as described below.

### Western blotting of heterologous TBE protein expression in *E. coli*

Protein expression in *E. coli* was confirmed by Western analysis, which was performed essentially as previously described (Walker *et al*., 2023) with minor modifications. *E. coli* overnight cultures expressing FZZ- or MBP-tagged proteins were diluted 1:100 in LB supplemented with 100 µg/ml carbenicillin and grown at 37 °C to OD_600_=0.4. Protein expression was induced with 0.1 mM IPTG, and cultures were transferred to 16 °C and grown for 18 h. The equivalent of 1 ml of OD_600_=1.5 culture was pelleted by centrifugation (12,000 x g for 5 min at 4 °C), and the pellet was suspended in 150 µl lysis buffer (20 mM Tris-Cl [pH 7.5], 150 mM NaCl, 0.5 mg/ml lysozyme). Samples were incubated at 25 °C for 10 mins before adding 10 mM DTT, 1% N-lauroyl sarcosine and 2X SDS loading dye (100 mM Tris-Cl [pH 6.8), 4% (w/v) SDS, 30 % (v/v) glycerol) and 0.07% (w/v) bromophenol blue). Samples were incubated at 95 °C for 10 min, and 5 µl of each sample underwent separation on a 4-12% gradient SDS-PAGE gel (Bio-Rad, Mini-PROTEIN TGX) at 100 V for 90 min. Proteins were transferred to a PVDF membrane (Invitrogen iBlot 2 Transfer Stack). Subsequent washing, blocking, and antibody incubation were performed by gentle nutation at room temperature. Membranes were washed with 1× PBS + 0.1% Tween 20, then incubated for 1 h in blocking buffer (1× PBS, 0.1% Tween 20, 5% BSA), washed in 1× PBS + 0.1% Tween 20, then incubated in primary antibody diluted in blocking buffer for 1 h at room temperature. For the FZZ-tagged samples, 1:40,000 monoclonal ANTI-FLAG M2 antibody produced in mouse (Sigma-Aldrich, F1804) was used. For the MBP-tagged sample, 1:10,000 of anti-TBE transposase antibody (GenScript) was used. For the loading control, 1:5,000 anti-GAPDH monoclonal antibody produced in mouse (Invitrogen, MA5-15738) was used. Membranes were washed three times in 1× PBS + 0.1% Tween 20, and primary antibodies were detected with either goat anti-mouse IgG1 heavy chain HRP (Abcam, ab97240) or goat anti-rabbit IgG (H+L)-HRP (Bio-Rad, 1706515). Finally, we performed enhanced chemiluminescence (ThermoFisher SuperSignal West Dura Extended Duration Substrate) and imaged blots on an Amersham Imager 600 (GE).

### Western blotting of native TBE protein expression in *Oxytricha*

∼300,000 mating cells from different time points were collected by centrifugation at 130 *g* for 2 min using a swinging bucket centrifuge (Sorvall RC6, Thermo Scientific). The supernatant was discarded, and the cell pellet was suspended in the residual medium. Next, 2x SDS loading buffer was directly added to the samples, which were then heated at 95 °C for 10 min. The samples were run on TGX Stain-Free Precast Gels (Bio-Rad) and transferred to a 0.2 µm Immun-Blot PVDF Membrane (Bio-Rad, 1620177) under semi-dry conditions using Trans-Blot SD blotter. The membrane was blocked with 5% nonfat dry milk (biokemix, M0841) in 1x TBS-T for 1 hour at RT and then incubated overnight at 4 °C on a rotator with a custom-made rabbit anti-TBE transposase antibody (1:1000 dilution in blocker). The following day, the membrane was washed five times with 1x TBS-T (5 min each) and incubated with 1:10,000 goat anti-rabbit IgG (H + L)-HRP (Bio-Rad, 1706515) for 1 hour at RT. After final washes (five times with 1x TBS-T, 5 min each), the chemiluminescence signal was detected using Clarity Western ECL reagent (Bio-Rad), and the membrane was imaged with a ChemiDoc imager (Bio-Rad).

### Chromatin-immunoprecipitation followed by next-generation sequencing (ChIP-seq) in *E. coli*

ChIP-seq in *E. coli* was performed essentially as described (Bonocora & Wade, 2015; Hoffmann *et al*., 2022; Park, 2009), with some modifications. Cultures were prepared and protein expression was induced as described in the Western blot method, above. After growing induced cultures at 16 °C for 18 h, formaldehyde was added to a final concentration of 1% to each 40 ml of culture and nutated for 20 min at room temperature. Formaldehyde was quenched by adding 4.6 ml of 2.5 M glycine, nutating for 10 min. Samples were then centrifuged at 4000 x g at 4 °C for 8 min. The cell pellet was washed twice in cold 1× TBS, and then the equivalent of 40 ml of OD_600_=0.6 was transferred to a tube to normalize cell density across samples and pelleted again by centrifugation. The pellet was transferred to a 1.5 ml Eppendorf tube and centrifuged at 10,000 x g at 4 °C for 5 min. The supernatant was discarded and cell pellets were flash-frozen in liquid nitrogen.

Cell pellets were resuspended in lysis buffer (50 mM HEPES-KOH [pH 7.5], 0.1% (w/v) sodium deoxycholate, 0.1% (w/v) SDS, 1 mM EDTA, 1% (v/v) Triton X-100, 150mM NaCl, 1× cOmplete Roche protease inhibitor), and transferred to a 1 ml milliTUBE AFA Fiber (Covaris). Samples were sonicated on an M220 Focused-ultrasonicator (Covaris) with the following SonoLab 7.2 settings: minimum temperature, 4 °C; set point, 6 °C; maximum temperature, 8 °C; peak power, 75.0; duty factor, 10; cycles/bursts, 200; sonication time, 17.5 min. After sonication, 10 µl of sheared cleared lysate was transferred to a separate tube as the pre-IP control (“input”). The remainder was used for immunoprecipitation.

For immunoprecipitation, 25 µl Dynabeads Protein G (Thermo Fisher Scientific) for each sample was washed four times with 1× PBS + 0.5% BSA and mixed with 4 µl monoclonal anti-FLAG M2 antibody produced in mouse (Sigma-Aldrich F1804), which was conjugated at 4 °C for 6 h. After conjugation, beads were washed four times with 1× PBS + 0.5% BSA, resuspended in 30 µl lysis buffer, then mixed with sonicated samples and rotated overnight at 4 °C.

The following day, beads were washed three times with lysis buffer lacking protease inhibitor, then washed once with lysis buffer lacking protease inhibitor and supplemented with 500 mM NaCl, once with ChIP wash buffer (10 mM Tris–HCl [pH 8.0], 250 mM LiCl, 0.5% (w/v) sodium deoxycholate, 0.5% (v/v) Nonidet-P40, 1 mM EDTA) and finally twice with TE buffer (10 mM Tris-HCl [pH 8.0], 1 mM EDTA). After the final wash, beads were suspended in 200 µl elution buffer (1% (w/v) SDS, 0.1 M NaHCO_3_) and incubated at 65 °C for 1 h 15 min to release protein-DNA complexes from the beads, vortexing every 15 min to resuspend the beads. While incubating the immunoprecipitated samples, 10 µl of the frozen pre-IP samples were withdrawn and 190 µl elution buffer was added. After the 65 °C incubation was complete, 10 µl of 5 M NaCl was added to each pre-IP and IP sample, which were incubated at 65 °C overnight to reverse crosslinks.

After overnight incubation, RNA was digested by adding 1 µl RNase A (Thermo Fisher Scientific) and incubating at 37 °C for 1 h. Protein was digested by adding 2.8 µl of 20 mg/ml proteinase K (Thermo Fisher Scientific) and incubating at 55 °C for 1 h. DNA was purified (QIAquick PCR Purification Kit) and eluted in 40 µl TE buffer before the concentration was measured by fluorometry (DeNovix dsDNA Ultra High Sensitivity Kit). Samples were normalized to the lowest concentration, and DNA was prepared for next-generation sequencing using the NEBNext Ultra II DNA Library Prep Kit for Illumina (NEB). ChIP-seq samples were sequenced on a paired-end run using NextSeq Mid or High Output Kits with 150 cycles (Illumina).

Paired-end reads were filtered, trimmed, and mapped as described in the Smear-seq workflow above. After read mapping, peaks were called using MACS2 (Zhang *et al*., 2008), filtered to exclude peaks with a score less than 20, and *de novo* motif prediction was performed using Homer using the command ‘findMotifsGenome.pl’ with default settings and a window size of 100 (Heinz *et al*., 2010).

### ChIP-seq in *Oxytricha*

*Oxytricha* cell lysates were prepared for ChIP-seq at 0 h, 36 h, and 48 h after mixing of cells of compatible mating type. Two million cells were fixed in 1% formaldehyde (Sigma) for 2 min, and crosslinking was quenched by adding glycine to a final concentration of 125 mM, followed by incubation for 5 min at RT on a rotator. Fixed cells were then lysed in 1 ml of ChIP lysis buffer (50 mM HEPES/KOH pH 7.5, 140 mM NaCl, 1 mM EDTA, 1% Triton X-100, 1% SDS, 0.1% sodium deoxycholate, 1x Protease Inhibitors Cocktail, Sigma) for 15 min on ice. Chromatin was sheared using Q800R3 Sonicator with 25 cycles of 30 seconds on/off with 40% amplitude, spun down, and the cleared lysate was flash frozen at −80 °C.

Each 1 ml sample was diluted 10x in dilution buffer (50 mM Tris-HCl pH 8.0, 150 mM NaCl, 1mM EDTA pH 8.0, 1% Triton, 1x cOmplete Roche Protease Inhibitor), and pre-cleared in Dynabeads Protein G. To bind the anti-TBE transposase antibody to the beads, 50 µl Dynabeads Protein G for each sample was washed two times with 1× PBST and mixed with 17.5 µl of custom anti-TBE transposase antibody produced in rabbits, which was conjugated at 4 °C for 6 h. After conjugation, beads were washed once with 1× PBST, resuspended in 50 µl dilution buffer, then mixed with the sonicated pre-cleared samples and rotated overnight at 4 °C. The following day, the remaining ChIP-seq steps (protein/RNA digestion, DNA cleanup, and NGS library preparation) were performed as described above, in the methods section for *E. coli* ChIP-seq.

### Indirect immunofluorescence

Immunofluorescent staining experiments were performed as previously described (Fang *et al*., 2012) with minor modifications. Briefly, cells were fixed with 4% paraformaldehyde (Electron Microscopy Sciences) for 10 min at RT on an end-to-end rotator, followed by two washes with 1x PBS. Fixed cells were placed on a Poly-L-Lysine-coated (0.1 mg/ml, Gibco) hydrophobic printed 12-well slide (Epredia, ER202W) and incubated overnight at 4 °C. The cells were then permeabilized with permeabilization solution (0.5% Triton X-100 in 1x PBS) for 30 min at RT, followed by a 5 min incubation in 0.1 M HCl. After a brief wash with 1x PBS, Image-iT™ FX Signal Enhancer (Invitrogen) was applied, and the cells were incubated for 30 min at RT. Cells were subsequently blocked with blocking buffer (0.2% Triton X-100, 0.5% IgG, and protease-free BSA in 1x PBS) for 1 hour at RT. Rabbit anti-TBE transposase antibody, diluted 1:100 in blocking buffer, was applied to the cells, and the slide was incubated overnight at 4 °C. Next day, the cells were washed three times with washing solution (0.2% Triton X-100 in 1x PBS) for 10 min each at RT. Alexa Fluor™ 488-conjugated goat anti-rabbit secondary antibody (1:1000 in blocking solution; Invitrogen, A11034) was then applied, and the slide was incubated at 37 °C for 1 hour in the dark. After a final series of three washes with washing solution (10 min each at RT), DAPI-containing VECTASHIELD® Antifade Mounting Medium (Vector laboratories, H-1000-10) was added to the samples, and the slide was covered with a long cover glass (Fisherbrand, 12545J). Fluorescent images were captured using a Zeiss Axiovert 200 inverted microscope equipped with an Axiocam 506 monochrome camera. Images were pseudo-colored using ImageJ64 when merging two channels.

## RESULTS

### Expression of the TBE transposase ORF induces widespread DNA fragmentation in *E. coli*

To study the function of a high copy number transposase implicated in genome rearrangements in *Oxytricha*, we set out to heterologously express candidate TBE transposases in *E. coli*, focusing our initial efforts on a representative transposon encoded in the *Oxytricha* MIC genome and referred to as TBE2.1(7905), a member of the TBE2.1 family encoded on contig 718000006<u>7905</u> (Chen *et al*., 2014). When we expressed the protein in *E. coli* BL21(DE3) and isolated and analyzed the cellular DNA fraction (**Figure 1B**), we observed a striking DNA fragmentation phenotype, characterized by a significant and reproducible DNA smear upon electrophoretic separation (**Figure 1C**). Apparent DNA degradation was visible as early as 3 hours after induction, and by 18 h, the DNA products ranged broadly in length. To determine whether this phenotype was merely a cytotoxic byproduct of heterologous protein over-expression, we cloned inactivated variants of the transposase, mutated in the predicted active site from DDE to D<u>A</u>E. This and other single amino acid alanine substitutions in the DDE motif abolished the DNA smearing phenotype (**Figure 1D**), providing the first direct evidence of TBE transposase-mediated DNA cleavage activity. Of note, these experiments in *E. coli* lacked any presence of bona fide TBE transposon DNA substrates.

We next set out to determine the composition of DNA products that appeared after transposase induction, via a high-throughput sequencing approach here dubbed ‘Smear-seq’ (**Figure 2A** and Materials and Methods). To prepare samples, we induced the WT or catalytically-inactivated transposase for either 1 h or 18 h, extracted DNA by miniprep, and then performed 3’ polyadenylation and second strand synthesis, followed by ligation to sequencing adaptors, and sequencing on an Illumina platform. After mapping sequencing reads to the TBE expression vector and the *E. coli* BL21(DE3) genome, we quantified the relative composition of plasmid or genomic DNA in each sample and found that after 18 h of induction, the WT transposase samples primarily comprised genomic DNA (>95% of reads), whereas the 3 h timepoint and the catalytically-inactivated transposase samples primarily comprised plasmid DNA (**Figure 2B**). These data suggest that presence of the TBE transposase active site permits extensive genomic DNA cleavage by the transposase in *E. coli*.

To test whether genomic DNA fragmentation was accompanied by cell death, we measured cell density by spectrophotometry and observed a dramatic loss of cell density after induction of the WT transposase, but not after induction of catalytically-inactivate transposase (**Figure 2C**), consistent with the genomic DNA cleavage activity of the wildtype TBE enzyme.

Our deep-sequencing approach also allowed us to search for preferred DNA cleavage sites, by examining the distribution of reads across the *E. coli* reference genome. We observed reproducible features in the WT 18 h sample on megabase-scale genomic regions within the *E. coli* genome, with consistent regions of enrichment and depletion and coverage profiles that were nearly indistinguishable across two independent biological replicates (**Figure 2D**). These data suggests that the TBE transposase may be preferentially cleaving consistent regions of the genome.

### ChIP-seq reveals degenerate DNA sequence motifs bound by TBE transposases

To more directly and globally profile specific DNA sites in the *E. coli* genome recognized by the TBE transposase, we applied ChIP-seq (chromatin immunoprecipitation followed by DNA sequencing) using the WT or catalytically-inactivated (DAE) transposase fused to a C-terminal epitope tag (3xFLAG) (**Figure 3A**), reasoning that the DAE variant would prevent DNA degradation and thus reveal the underlying substrate specificity. Examining the ChIP-seq read coverage across the *E. coli* genome revealed narrow regions of enrichment and depletion that were highly consistent across biological replicates (**Figure 3B**), in stark contrast to the more diffuse Smear-seq signal (**Figure 2D**). We induced each transposase variant for 3 h and 18 h and observed strong agreement between the ChIP-seq profiles for all samples excluding WT at 18 h, which had more diminished peaks, we conjecture due to accumulated DNA cleavage activity reducing the signal.

**Figure 3.**
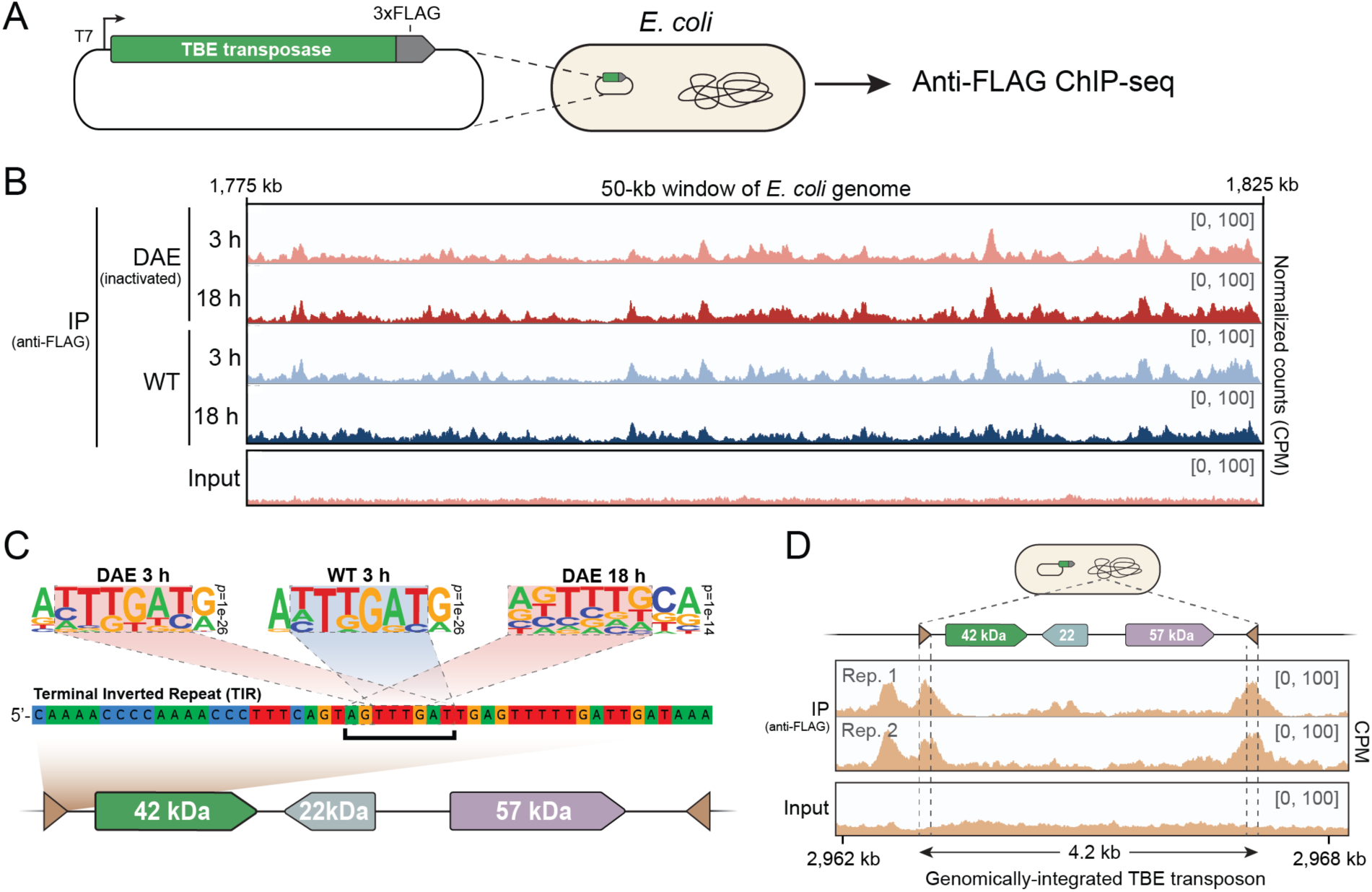
ChIP-seq reveals DNA sequence preferences for degenerate motifs partially encoded within the TBE transposon ends. (**A**) ChIP-seq approach for determining DNA binding preferences of the TBE transposase in *E. coli.* (**B**) Read coverage across a representative 50-kb window of the *E. coli* genome, for the WT and catalytically-inactivated transposase (DAE). The y-axis represents counts per million (CPM) and the axis limit is set to 100. (**C**) Sequence logos represent the top statistically significant motif from *de novo* motif prediction (Homer; Heinz *et al*., 2010). P-values indicate statistical significance; motifs are considered significant if their p-values are less than 1e-10. Motif prediction for the WT 18 h sample did not detect any statistically significant motifs (N.S.). Dashed boxes indicate regions that map to the TBE transposon ends, as illustrated by the first 50 nt of the TBE2.1 terminal inverted repeat (TIR). (**D**) Read coverage from ChIP-seq using a catalytically-inactive TBE2.1 transposase induced for 18 h in a strain harboring a genomically-integrated TBE2.1 transposon. Vertical dashed lines on the coverage plots indicate the TIR boundaries.

To test whether conserved DNA sequence features were present within the genomic regions of enrichment, we performed peak-calling and *de novo* motif prediction. This analysis revealed a short sequence motif (5’-ANTTTGA) as the top motif detected in all samples except for WT 18 h, where no statistically significant motifs were found (**Figure 3C**). This motif was not only called as the top motif in each sample, but also the only motif that met the significance threshold in each of the three samples (**Figure S1A**). The absence of statistically significant motifs in the WT 18 h sample is likely also due to transposase cleavage activity leading to cleavage and product release, thus decreasing enrichment at preferred binding sites. Intriguingly, the short sequence motif is also found within the TBE transposon terminal inverted repeats (**Figure 3C, TIR**), which transposases typically recognize to mobilize an element.

We hypothesized that this motif would resemble sequence features found within the TBE transposon terminal inverted repeat (TIR) ends, and that structural modeling might identity likely points of interaction between the transposase and TIRs. To test this, we used AlphaFold Multimer (Jumper *et al*., 2021) to predict the structure of the TBE transposase dimer, which suggested that it was similar to the solved structure of another Tc*1*/*mariner* family transposase, Mos1, with an RMSD of 7.991Å. (**Figure S1B**). Mos1, first isolated from *Drosophila mauritiana* (G. Bryan *et al*., 1990; G. J. Bryan *et al*., 1987), encodes an N-terminal DNA binding domain (residues 1-112) and C-terminal DD[D/E]-family catalytic domain (residues 126-345). Each N-terminal DNA binding domain encodes two ⍺-helical motifs (HTH1 and HTH2) which contact the *mariner* transposon ends at positions 21-26 and 8-13, respectively (Richardson *et al*., 2009). Intriguingly, when we predicted the structure of the TBE transposase with 40 bp of TIR DNA (RoseTTAFoldNA; Baek *et al*., 2023), its two HTH motifs were positioned near the TTT trinucleotide motifs at the transposon ends, with the DNA binding domains contacting TIR DNA at positions 24-27 and 16-18, respectively (**Figure S1C**). The TTT trinucleotide motif was also detected in the *E. coli* ChIP-seq peaks (**Figure 3C**), lending additional support to the interpretation that the transposase exhibits DNA sequence preferences for this motif.

Next, we set out to test whether we could detect transposase enrichment within the native TBE transposon end sequences. We generated a genomically-modified *E. coli* strain that harbored a complete TBE transposon element in its circular chromosome and performed ChIP-seq in this new genetic background (**Figure 3D**). We observed strong enrichment within the transposon terminal inverted repeats (TIRs), offering the first evidence that the TBE transposase recognizes its transposon ends and suggesting that the TBE2.1 transposase may be actively targeting TBE elements during *Oxytricha* genome rearrangement.

To test whether the transposase could also recognize *Oxytricha* genomic elements that are eliminated during programmed DNA rearrangement, we cloned a ∼4.5-kb region of micronuclear DNA covering the TEBPβ gene locus, which has the most thoroughly studied rearrangement products (Nowacki *et al*., 2008, 2009). To avoid reintegrating the *Oxytricha* genomic element into the *E. coli* genome, we first tested whether we could recapitulate ChIP-seq results with a plasmid-encoded TBE transposon element, and indeed observed consistent enrichment profiles between the genomic- and plasmid-encoded elements (**Figure S2A**). When we examined the enrichment profile across the native TEBPβ locus, introduced on a bacterial plasmid, we observed a variable pattern of enrichment clustered near eliminated regions (**Figure S3B**), with read coverage highest in a region with a high-density of rearrangement junctions and IESs. This result may indicate that other protein or nucleic acid features are responsible for targeting transposase activity to IESs during rearrangement.

### TBE accessory proteins do not alter TBE DNA binding profiles in *E. coli*

The TBE transposon encodes two proteins in addition to the putative transposase (**Figure 1B**), which are under purifying selection and are proposed to function as accessory factors during transposition (X. Chen & Landweber, 2016). This feature is surprising for Tc*1*/*mariner* elements, which typically encode only their transposase and generally lack accessory proteins (Dupeyron *et al*., 2020). One of the two *Oxytricha* proteins is estimated to be 57-kDa and comprises predicted zinc-finger and kinase domains, while the other is estimated to be 22-kDa protein with no predicted domains (X. Chen & Landweber, 2016) (**Figure S3A**). To assay in *E. coli* whether the expression of these accessory proteins could affect DNA cleavage or binding of the TBE transposase, we prepared epitope-tagged expression vectors for each component and co-expressed them with untagged dual-expression vectors for the other components (**Figure S3B**). We first confirmed protein expression of each element by Western blot and observed bands at the expected size for each protein, as well as a second, smaller band for the 57-kDa sample that might represent a protein degradation product (**Figure S3C**). ChIP-seq against tagged versions of either accessory protein or the TBE transposase, in the presence or absence of the two other untagged proteins, did not display differences in the DNA-binding profiles for any combination relative to the pre-IP input profile, suggesting that the accessory proteins might not bind DNA with specificity (**Figure S3D**) or that the *E. coli* expressed versions might not capture the *Oxytricha* environment or modifications necessary for function. It is also possible that the accessory proteins participate in later steps of transposition or repair during genome rearrangement.

### Transposase homologs exhibit similar DNA sequence preferences

Phylogenetic analysis of the 42-kDa transposase ORF encoded within diverse TBE transposons revealed that they cluster into four families — TBE1, TBE2.1, TBE2.2, and TBE3 (**Figure 4A**) (X. Chen & Landweber, 2016) — and allowed the generation of consensus protein sequences for each family that share >70% amino acid identity (X. Chen & Landweber, 2016) and possess conserved catalytic active sites (**Figure S4**). We wondered whether differences among transposase families might confer differences in transposase activity and therefore repeated the DNA fragmentation and ChIP-seq assays using the consensus sequences for each of the four TBE families (**Figure 4B-D**). Each transposase representative induced a similar DNA fragmentation phenotype, although the phenotype was less dramatic for the TBE3 family sequence (**Figure 4B**). We also prepared catalytically-inactive variants of each transposase family sequence and confirmed that DNA fragmentation required the presence of an intact catalytic triad. ChIP-seq experiments using the consensus sequences revealed consistent profiles of DNA enrichment across the *E. coli* genome, although the profiles for the TBE1 and TBE3 representatives were less pronounced (**Figure 4C**). The most noticeable difference when using consensus sequences arose when we performed ChIP-seq in *E. coli* strains encoding genomically-integrated TBE transposons from each family. In these genomic backgrounds, we did not detect strong enrichment within the transposon TIRs (**Figure 4D**). This result could be attributed to differences in the DNA binding preference, or to differences between the transposon TIRs among families, which differ up to 47% (X. Chen & Landweber, 2016).

**Figure 4.**
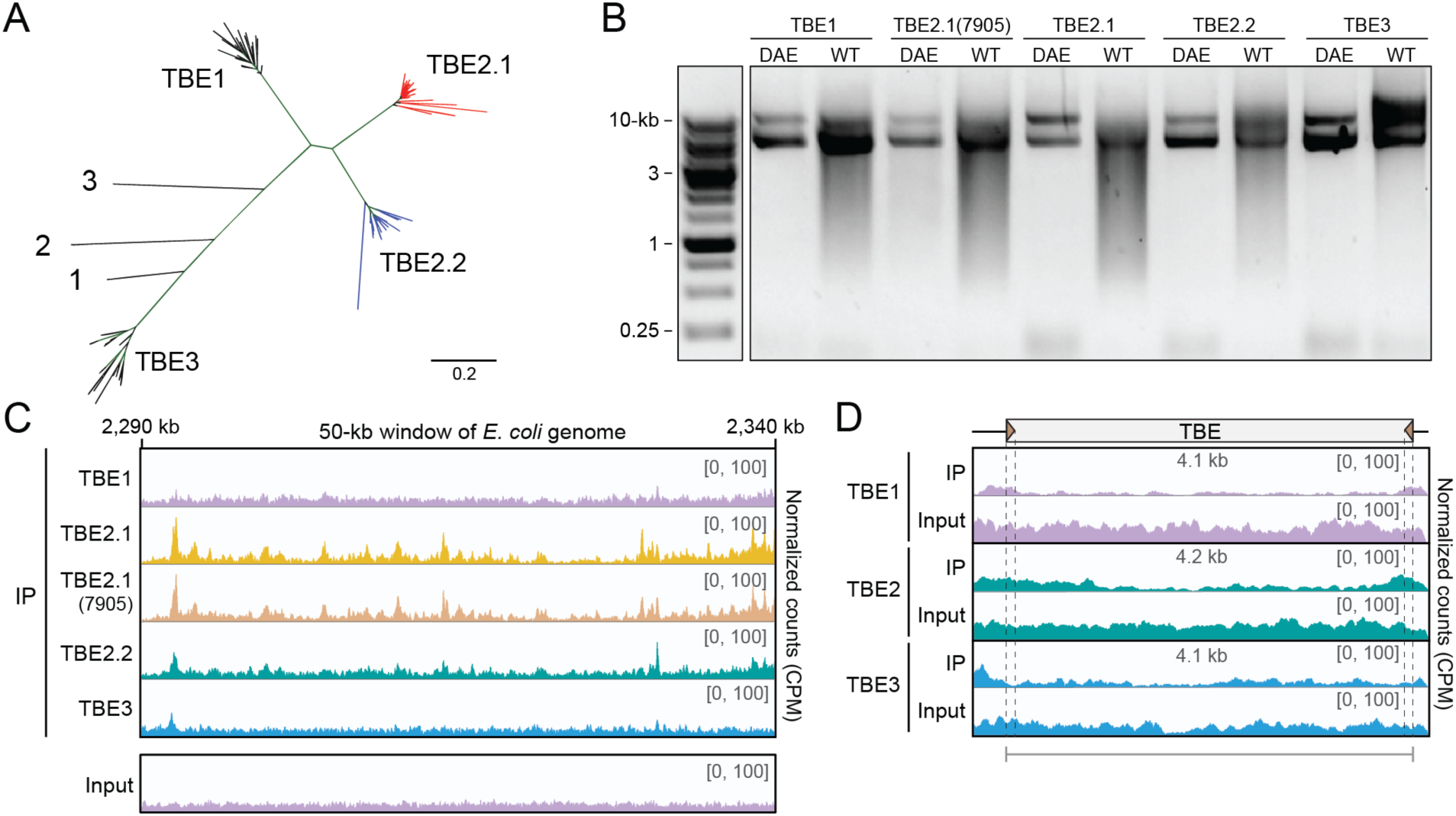
TBE transposase homologs are catalytically active and exhibit similar enrichment profiles across the *E. coli* genome. (**A**) Unrooted Bayesian phylogenetic tree of four families of TBE elements, from (X. Chen & Landweber, 2016), built using the protein sequences of the 57 kDa protein. The scale indicates branch substitutions per site, and branches labeled 1-3 are the following TBE orthologs: 1: *Sterkiella histriomuscorum*; 2: *Tetramena sp.;* 3: *Laurentiella sp*. (**B**) DNA fragmentation assay for representative TBE transposase homologs from each of the four major families, upon heterologous expression in *E. coli*. Each sample was induced for 18 h before DNA was purified, and 500 ng DNA was separated by electrophoresis on a 1% agarose gel and visualized using SYBR Safe DNA Gel Stain. A catalytically inactivated mutant (DAE) was generated and cloned for a representative of each TBE family. For each transposase representative, only the WT transposase induces DNA fragmentation. (**C**) Read coverage across a representative 50-kb window of the *E. coli* genome from ChIP-seq using the catalytically-inactivated representative from each TBE group. The y-axis represents counts per million (CPM) and the axis limit is set to 100. (**D**) Read coverage in CPM from ChIP-seq in strains harboring a genomically-integrated TBE element corresponding to a representative from either TBE1 (from ctg7180000067530), TBE2 (from ctg7180000069065), or TBE3 (from ctg7180000089059). The y-axis represents CPM with the axis limit set to 100, and dashed lines indicate the TIR boundaries.

### Transposase binding correlates with DNA cleavage

We wondered whether the DNA signatures from our ChIP-seq and Smear-seq approaches overlapped, which would indicate correspondence between DNA binding and cleavage activity. Visual inspection of the read coverage profiles from both datasets revealed an apparent consistency between the ChIP-seq and 18 h WT Smear-seq samples, despite the more diffuse mapping in the Smear-seq profiles compared to the ChIP-seq profiles, which could be due to downstream DNA degradation after transposase-mediated DNA cleavage (**Figure 5A**, **Figure S5A**). To test whether this qualitative similarity was statistically significant, we averaged the genome-wide read coverage within 2-kb bins and compared the average coverage between samples (**Figure 5B-D**, **Figure S5B**). This analysis revealed a moderate positive correlation (*r =* 0.48) between the ChIP-seq signal and the Smear-seq signal after 18 h of inducing the catalytically active transposase (**Figure 5C**). The moderate correlation disappeared when we compared the ChIP-seq signal to the Smear-seq signal after inducing the catalytically-inactive transposase (**Figure 5D**). We also observed a remarkable consistency between Smear-seq replicates at 18 hours using the WT transposase (*r* = 0.99 **Figure S5B**), reflecting reproducible and thorough DNA degradation in the presence of catalytically active *Oxytricha* transposase expressed in *E. coli*, confirming efficient expression. The correlation patterns remained consistent across all bin sizes ranging from 500 bp to 10 kb (**Figure S5C**). Overall, the consistency between regions of DNA binding, as measured by ChIP-seq, and regions of DNA cleavage, as measured by Smear-seq, indicates that the DNA recognition and cleavage activity of the TBE transposase is sequence-specific, favoring degenerate motifs that are partially present within the TBE transposon ends.

**Figure 5.**
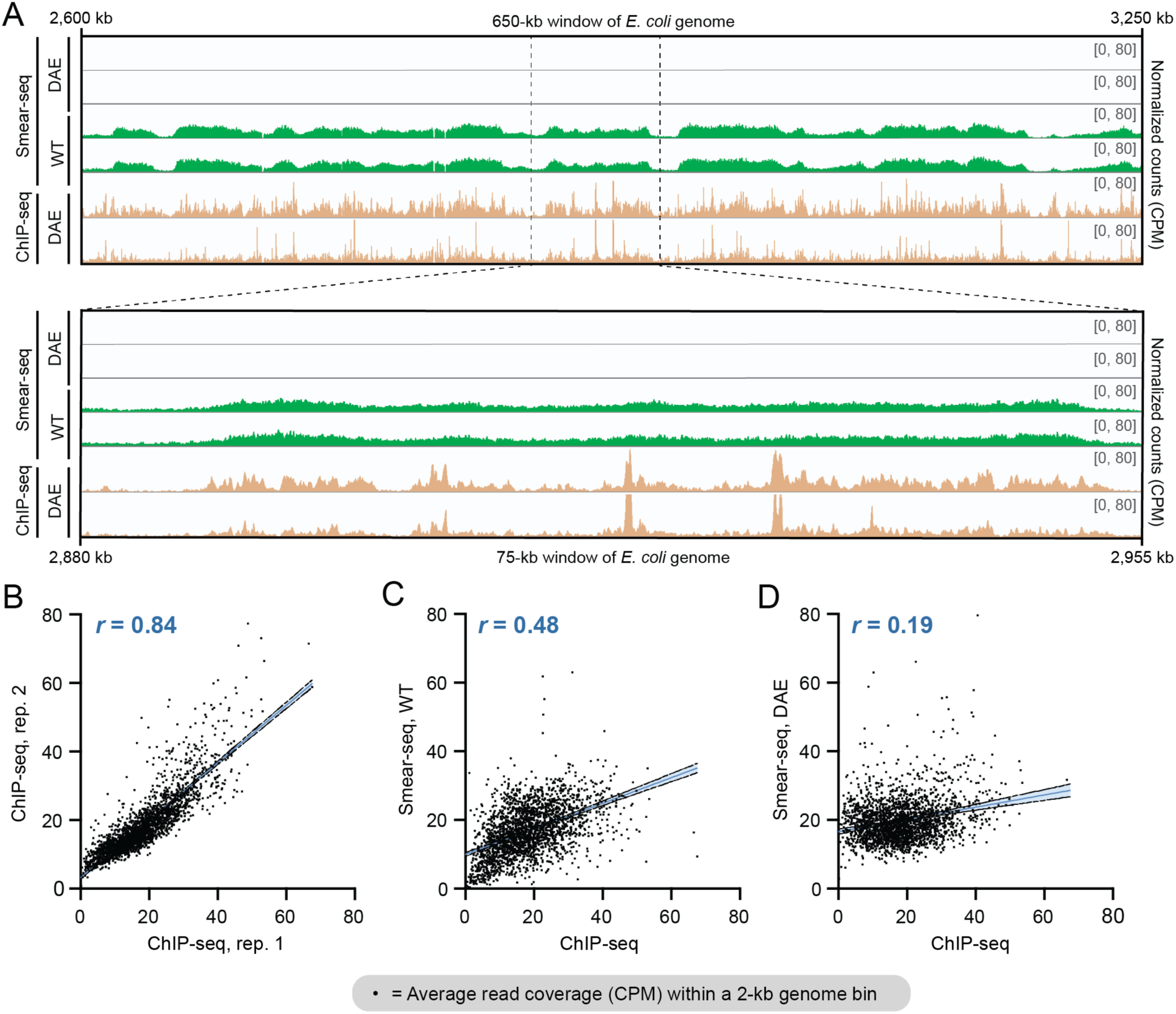
Transposase binding activity (ChIP-seq) correlates with cleavage activity (Smear-seq). (**A**) Read coverage profiles of the 18 h Smear-seq DAE and WT samples, compared to the 18 h ChIP-seq DAE signal, in 650-kb (top) or 80-kb (bottom) bins across the *E. coli* genome. The y-axis represents CPM with the axis limit set to 80. (**B-D**) Comparison between average read coverage (CPM) across 2-kb bins of the *E. coli* genome. There is strong correlation between ChIP-seq replicates (**B**), moderate correlation between ChIP-Seq DAE and 18 h WT Smear-seq samples (**C**), and no correlation between the ChIP-seq DAE and the 18 h DAE Smear-seq signal (**D**). *r* values represent the Pearson correlation, and linear regressions are shown with 95% confidence bands of the best-fit line.

### TBE transposase is expressed in the developing nucleus and preferentially associates with TBE elements in *Oxytricha*

Efforts to study the biochemical properties of TBE transposases in vitro have been hampered by protein insolubility (Orsolya Barabas, personal communication), so we decided to pursue alternative experiments to monitor its native localization and activity in *Oxytricha*. We selected a 14-amino-acid peptide corresponding to the predicted surface-exposed region between the two N-terminal DNA binding domains (**Figure S6A**), which is conserved in TBE2.1 and TBE2.2 (**Figure S6B**), and purified polyclonal antibodies after rabbit immunization. Western hybridization upon heterologous TBE transposase expression in *E. coli* revealed a clear band at the expected size (**Figure S6C**), supporting the specificity of the antibody.

Next, we performed a mating time course of *Oxytricha* cells and collected lysates at 12-hour intervals for anti-TBE transposase western analysis to monitor transposase expression. A strong 42-kDa band was present at 36 and 48 h, corresponding to the expected peak of genome rearrangement (**Figure 6A**) (X. Chen *et al*., 2014; Williams *et al*., 1993). To investigate TBE transposase localization, we performed immunofluorescence at the same 12-hour intervals post-mixing and observed a strong signal in developing somatic nuclei (**Figure 6B**). These data reveal that TBE transposases are not only expressed during development, but that they specifically localize to *Oxytricha*’s developing somatic macronucleus.

**Figure 6.**
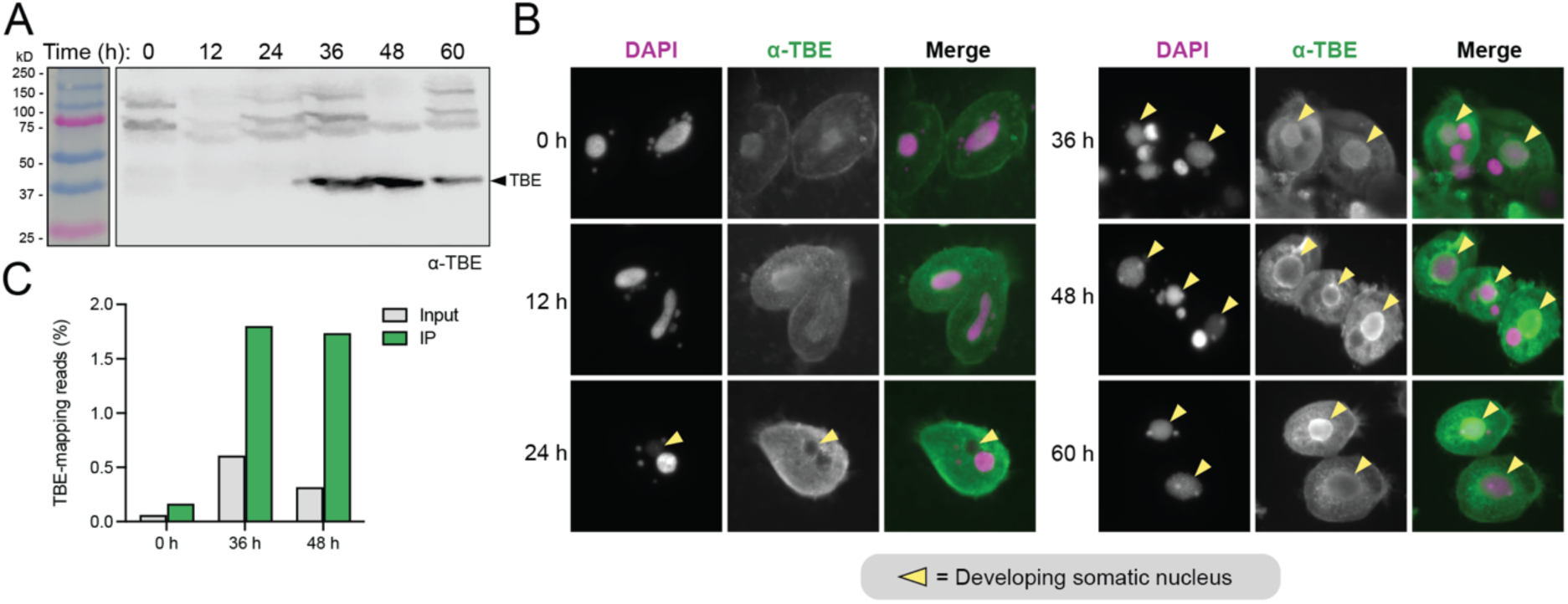
TBE transposase expression, localization, and transposon-DNA preference during *Oxytricha* development. (**A**) Western blot analysis using an anti-TBE transposase antibody against *Oxytricha* cell lysates collected at 12-hour intervals post mixing of compatible mating types. (**B**) DAPI staining of DNA and anti-TBE transposase immunostaining of *Oxytricha* cells at 12-hour intervals post mixing. Arrowheads indicate developing somatic macronuclei. Note that until 48 h, DAPI stains the parental macronuclei more strongly than the new macronuclei, due to the copy number difference. (**C**) Percent of micronuclear genome-mapping ChIP-seq reads that map to TBE2.1(7905) in the input and IP samples, at 0 h and two timepoints after mixing.

Finally, we performed ChIP-seq experiments in *Oxytricha* cells at 0, 36, and 48 hours post-mixing. When we mapped sequence reads back to the TBE-family transposons and compared the proportion of reads mapping to TBEs versus to the remainder of the micronuclear genome, we observed a ∼4-fold enrichment compared to input, and a striking >8-fold enrichment of TBEs at developmental timepoints compared to the 0 h timepoint (**Figure 6C**), revealing that TBE transposases preferentially target TBE transposon elements natively in *Oxytricha*. We did not observe peaks of enrichment within the transposon end sequences themselves (**Figure S6D**), which may be due to read-mapping artifacts; because we are mapping ChIP-seq reads to a single TBE2.1(7905) element, the sequence variability flanking the high copy-number TBEs heterogeneously distributed throughout the genome may cause coverage to drop out near the boundary of the element. Taken together, these data reveal that *Oxytricha* TBE transposases are expressed in their developing somatic nucleus, and preferentially associate with TBE-specific DNA transposons.

## DISCUSSION

By leveraging a heterologous expression strategy in *E. coli*, our study provides the first insights into TBE transposase activity, revealing the enzyme’s DNA sequence preferences for binding and cleavage. Our heterologous expression approach addresses a critical gap in understanding the DNA substrate preferences of this enzyme that participates in genome rearrangement. Functional studies of the TBE transposase in *Oxytricha* have been limited by tools to manipulate a high copy element in the germline genome. Our present approach overcomes these limitations and provides insight into the mechanism of DNA targeting and cleavage. The approach may be leveraged to study the activity of other transposases, which can be notoriously difficult to purify due to their basic properties and aggregation propensities (Jaillet *et al*., 2014), limiting the ability to study their biochemical properties and activities *in vitro*.

After observing an unexpected DNA fragmentation pattern when we expressed the TBE transposase in *E. coli*, we developed a deep sequencing approach (‘Smear-seq’) to investigate the enzyme’s genomic DNA fragmentation activity. By leveraging an orthogonal approach to directly profile DNA binding sites of the catalytically inactive transposase using ChIP-seq and *de novo* motif predictions, we discovered that the transposase preferentially binds short sequence variations of the motif 5’-ANTTTGA, which is partially present within the TBE transposon ends. Moreover, ChIP-seq revealed enrichment of the transposase within the native TBE end sequences (TIRs) of the TBE2.1(7905) variant. We further applied our scalable approach to profile DNA binding specificities for all major families of TBE transposases, revealing their ability to induce DNA fragmentation and to display consistent enrichment profiles across the genome. Finally, we found a moderately positive correlation between the ChIP-seq and Smear-seq signals, indicating that the TBE transposase not only binds DNA at specific sites but can also induce DNA fragmentation at these loci.

One limitation of the heterologous workflow is that it does not reconstitute native protein interactions between the TBE transposase and other *Oxytricha* elements that are absent from the *E. coli* expression system. We also acknowledge that our experiments primarily examined a single TBE2.1 transposase variant and consensus sequences from the four transposon families, which cannot represent the activity of the >34,000 specific TBE elements encoded in the MIC genome. Importantly, the majority of germline-encoded transposase genes do not contain an intact catalytic triad (X. Chen & Landweber, 2016), indicating that they have lost DNA cleavage activity and could be pseudogenes. However, non-catalytic paralogs could also serve other functions as binding partners or interactors, as in another model ciliate, *Paramecium tetraurelia* (Bischerour *et al*., 2018).

Transposases involved in genome rearrangement have been extensively profiled in other model ciliates, which also undergo large-scale genome rearrangements, albeit with reduced complexity and at a smaller scale than *Oxytricha*. *Paramecium* genome rearrangement eliminates >45,000 IESs and requires a domesticated PiggyMac (PGM) transposase (Arnaiz *et al*., 2012; Baudry *et al*., 2009). PGM is homologous to the *piggyBac* transposase, originally isolated from a cabbage looper moth, and it inserts precisely and exclusively into TTAA target sites (Fraser *et al*., 1983). In *Paramecium*, PGM is expressed in the developing macronucleus shortly after mating, and knockdown of PGM reduces cell viability and inhibits IES excision (Baudry *et al*., 2009). Not all copies of PiggyMac are catalytically intact; five groups of catalytically-inactivated Pgm-like proteins (Pgmls) interact with PiggyMac, and are required for both nuclear localization and DNA elimination activity (Bischerour *et al*., 2018). Intriguingly, the DNA-targeting activity of the catalytically-intact PGM is influenced by chromatin state, and the nucleosome remodeling protein ISWI1 participates in IES nucleosome depletion, perhaps by increasing chromatin accessibility for DNA cleavage by PGM (Singh *et al*., 2022). Although our work in *Oxytricha* indicates that the TBE transposase is sequence-specific, we have yet to determine the extent to which chromatin state modulates transposase recruitment or cleavage activity.

The genome of another well-studied ciliate, *Tetrahymena*, also encodes domesticated *piggyBac*-like transposases that are implicated in DNA rearrangement. Three of these transposases, Tpb1, Tpb2, and Tpb6 are also expressed shortly after mating and their knockdown impairs viability and inhibits DNA elimination (Cheng *et al*., 2010, 2016). The targeting mechanism of Tpb2 is based on recruitment to heterochromatin structures, as opposed to specific sequences (Cheng *et al*., 2010), while Tpb1/6 are thought to require sequence recognition at the boundaries of eliminated sequences (Cheng *et al*., 2016). Future work is needed to determine whether DNA recruitment of the TBE transposase is dependent on heterochromatin structures in its native environment; however, the short length of many *Oxytricha* IESs presents a challenge to chromatin recognition (Chen *et al*. 2014).

The relaxed sequence specificity of the TBE transposases differs markedly from the strict sequence-specific activity of most transposases that exclusively target their transposon end sequences to minimize off-target cleavage activity that could compromise host fitness. Although our work demonstrates that the TBE transposases do recognize their end sequences, they also exhibit remarkable off-target activity, with the ability to reproducibly bind and cleave distinct regions across the *E. coli* genome. Future work in *Oxytricha* is needed to confirm whether this relaxed specificity is indeed critical to facilitate the extensive cleavage events that occur during development of the somatic macronucleus. Together, this work reveals DNA binding and cleavage activity of the TBE transposases involved in *Oxytricha* genome rearrangement, and describes a facile heterologous approach to profile the activity of enzymes implicated in DNA rearrangements.

## DATA AVAILABILITY

High-throughput sequencing data and custom scripts used for analyses of high-throughput sequencing data are available at Zenodo (DOI 10.5281/zenodo.15040686). Datasets generated and analyzed in the current study are available from the corresponding authors on request.

## SUPPLEMENTARY DATA

Supplementary Data are available online.

## ACKNOWLEDGEMENT

We thank Sheela George, Ryan Tran, Jeeva Abraham, Sanjana Pesari and Zara Akhtar for laboratory support. We thank Bill Jack, Rodney Rothstein, Eric Greene, and all current and past Landweber and Sternberg lab members for discussions about *Oxytricha* biology. We thank Michael Lu and Tanner Wiegand for bioinformatics advice. We thank Margarita Angelova for figure design inspiration for *Oxytricha* genome rearrangement. We thank George Lampe and the JP Sulzberger Columbia Genome Center for NGS support.

## CONTRIBUTIONS

M.W.G.W., T.A., S.H.S., and L.F.L. conceived and designed the project. M.W.G.W. and T.A. designed and constructed TBE protein expression vectors and performed DNA fragmentation assays in E. coli. The DNA fragmentation phenotype was originally identified by T.A. M.W.G.W. performed and analyzed the ChIP-seq and Smear-seq experiments and validated the TBE transposase antibody in *E. coli*. D.J.V. and T.A. designed the Smear-seq approach. E.A. performed immunofluorescence experiments and western blotting in *Oxytricha*, and prepared *Oxytricha* lysates for ChIP-seq. H.S.F. and D.J.V. assisted in the analysis of ChIP-seq data. H.L. assisted in the construction of expression vectors. M.W.G.W wrote the manuscript with assistance from S.H.S. and L.F.L and input from all authors.

## SUPPLEMENTARY FIGURES

**Figure S1.**
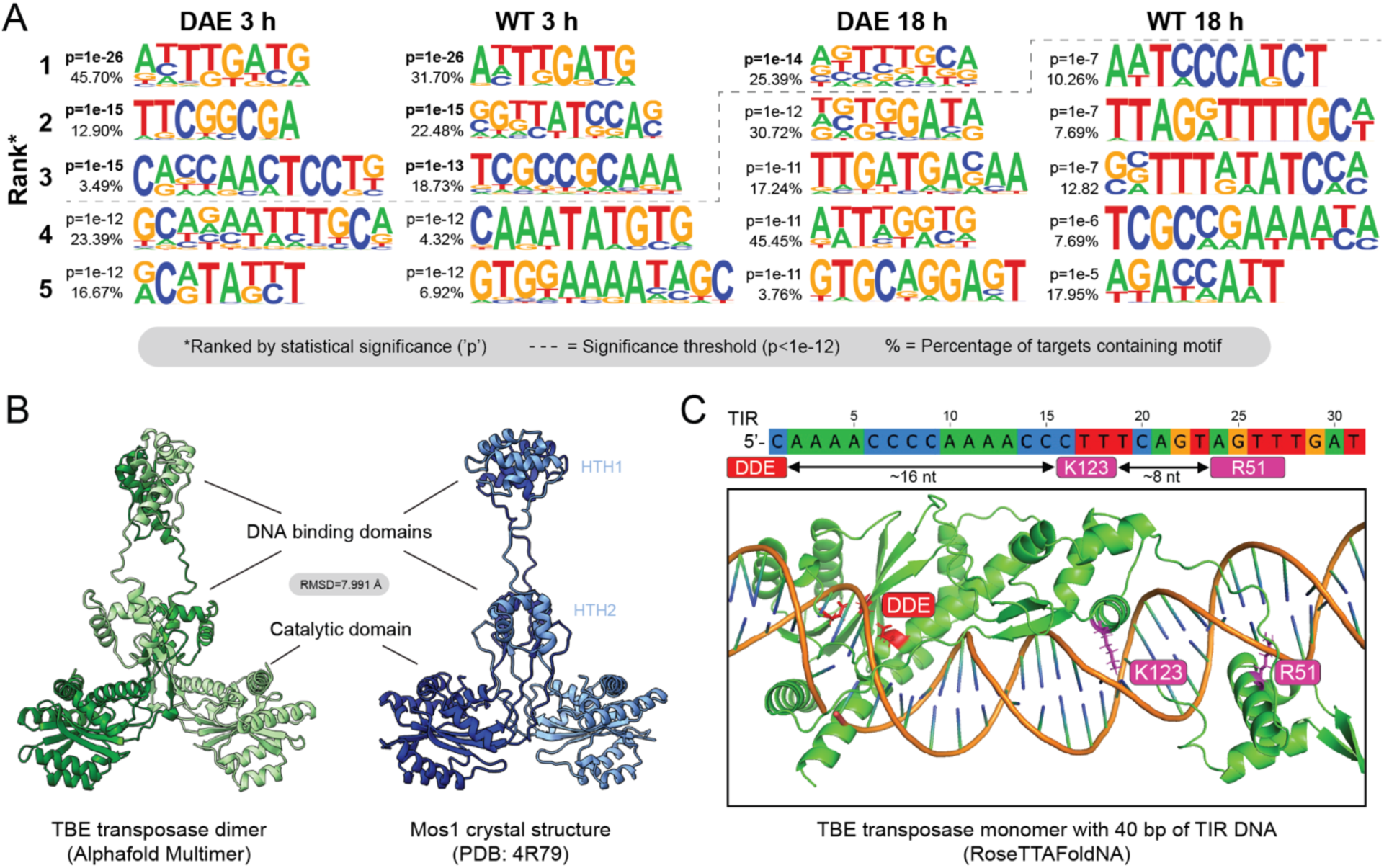
*De novo* motif analysis and structural prediction of the TBE transposase places putative DNA binding domains near preferred motifs within transposon ends. (**A**) Sequence logos identified from *de novo* motif prediction (Homer; Heinz *et al*., 2010) within peaks of ChIP-seq read enrichment for the catalytically-inactivated (DAE) and WT transposase. Sequence logos are ranked by statistical significance, and the top five most significant motifs from each sample are shown. The dashed line indicates the cutoff for statistical significance (p-value<1e-12). The percentage of ChIP-seq peaks containing each motif is listed to the left of each motif (%). (**B**) AlphaFold-Multimer (Evans *et al*., 2022) prediction of the TBE2.1(7905) transposase dimer (left). Annotated DNA binding domains are based on structural homology to the Mos1 crystal structure (right). DNA-binding helix-turn-helix (HTH) and catalytic domains are indicated. (**C**) RoseTTAFoldNA (Baek *et al*., 2023) prediction of a TBE transposase monomer interacting with TBE TIR DNA. The TIR DNA schematic (top) illustrates the predicted interaction sites of the DNA binding domain residues (K123 and R51) based on the RoseTTAFoldNA model.

**Figure S2.**
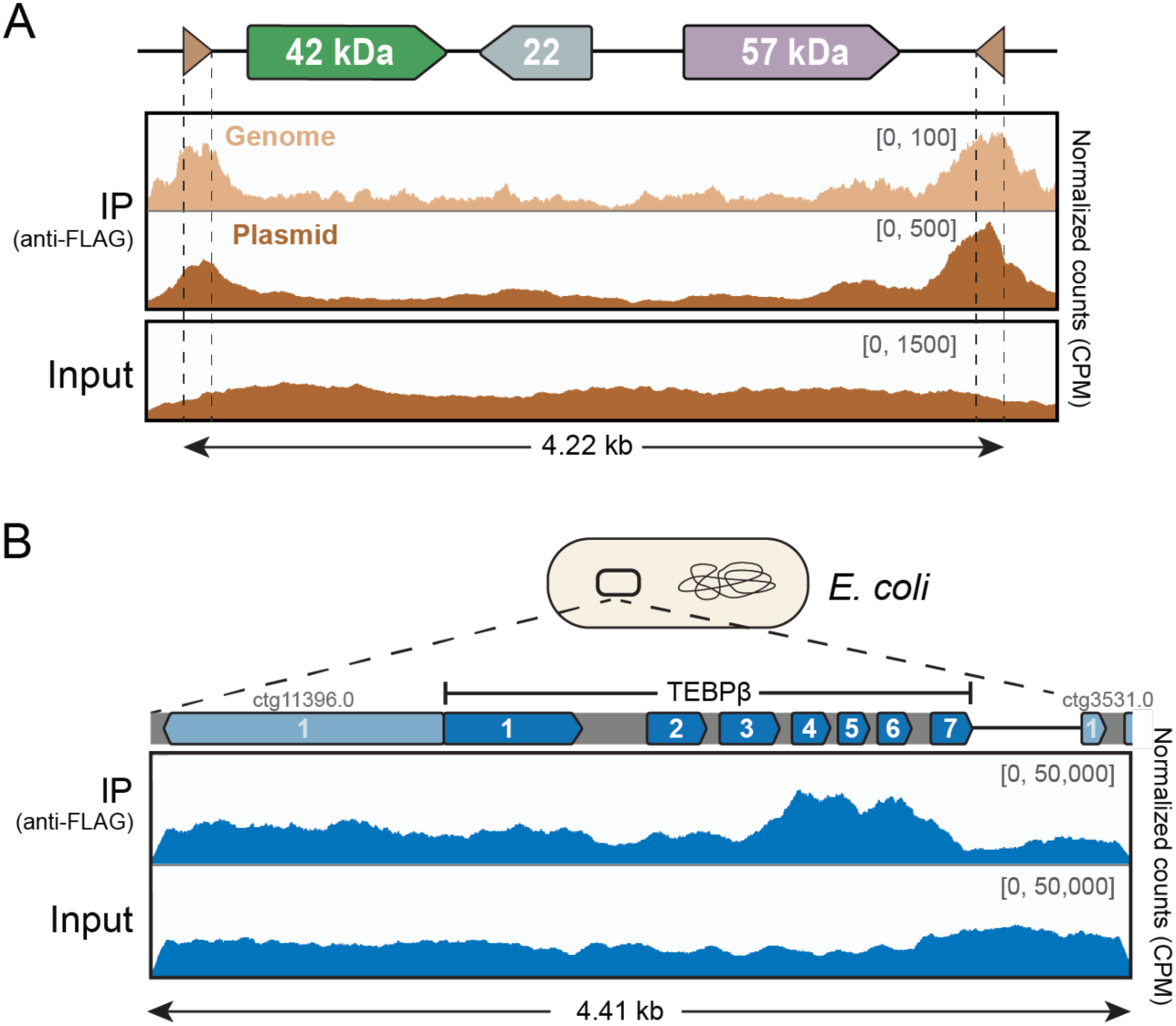
The TBE transposase does not exhibit a consistent pattern of enrichment across *Oxytricha* MIC genomic DNA in *E. coli*. (**A**) Read coverage from ChIP-seq of the TBE transposase using either a genomically-integrated (“genome”) or a plasmid-encoded complete TBE transposon (“plasmid”) in *E. coli*. The y-axis represents CPM and is set to the following maximum for each sample: 100 CPM for the genome IP track; 500 CPM for the plasmid IP track; and 1500 CPM for the plasmid input track. (**B**) ChIP-seq read coverage from the FZZ-tagged TBE transposase expressed in *E. coli* cells containing a cloned *Oxytricha* MIC genomic region on a plasmid containing the TEBPβ MIC locus and flanking regions. The y-axis represents CPM and is set to 50,000 for both the IP and input samples. Dark blue boxes within the TEBPβ MIC locus depict its 7 MDSs, gray boxes represent IESs, and pale blue boxes are MDSs for flanking loci; figure approximately to scale.

**Figure S3.**
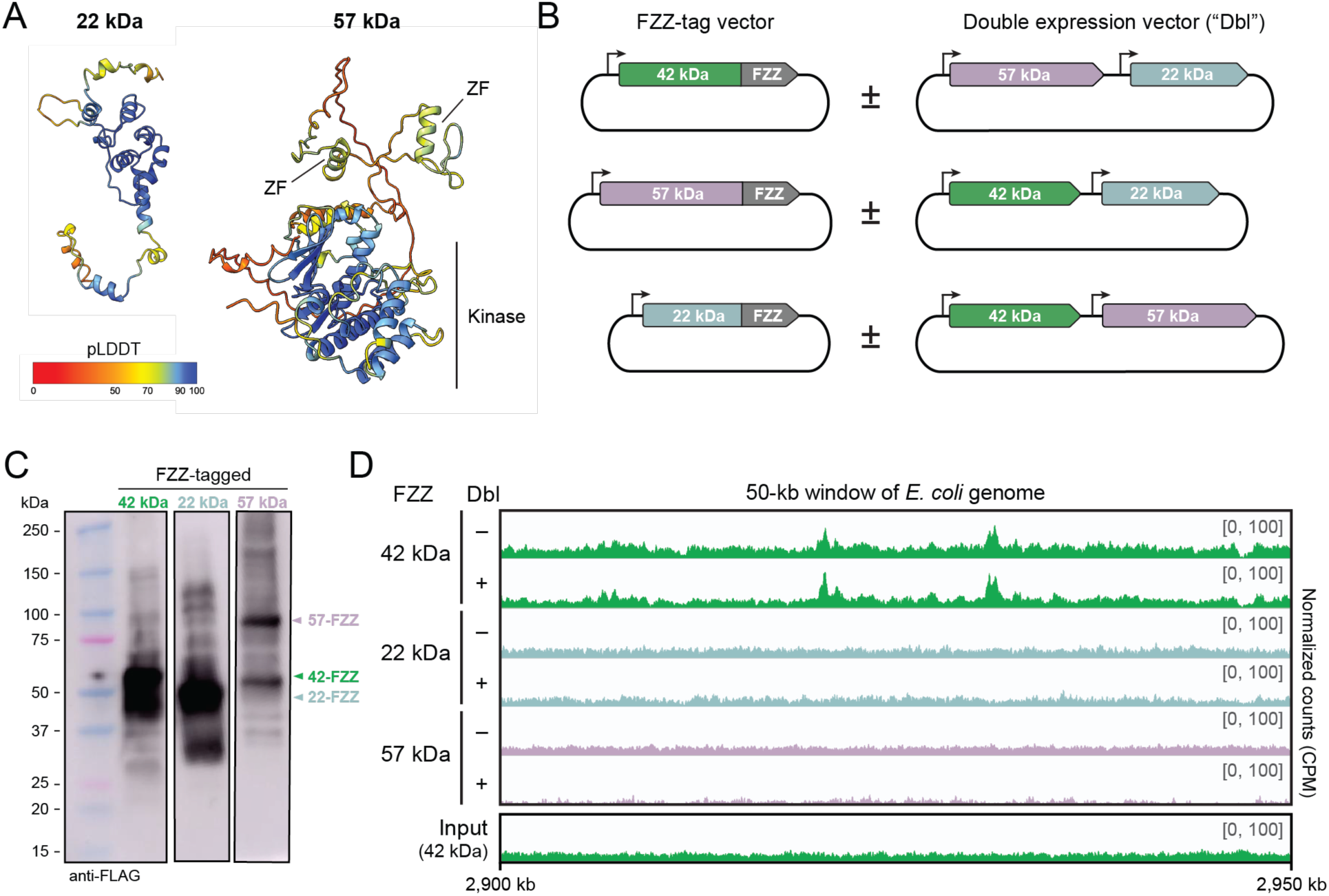
Investigating the accessory proteins encoded within TBE transposons. (**A**) AlphaFold predictions of the 22-kDa and 57-kDa ORF, colored by pLDDT (predicted local distance difference test) values. Predicted zinc finger (‘ZF’) and kinase domains from HHpred (Söding *et al*., 2005) are annotated on the 57-kDa structure; the 22-kDa protein exhibits no high-confidence domain predictions via HHpred. (**B**) Schematic of expression vector design for ChIP-seq experiments in D. In each experiment (horizontal rows in D), strains contained pairwise combinations of FZZ-tagged single-protein vectors in the presence and absence of untagged double protein expression vectors (‘Dbl’). (**C**) Western blot with an anti-FLAG antibody used against each of the three FZZ-tagged expression vectors in *E. coli*. Expected sizes for the FZZ-fusion proteins are 77 kDa (57-kDa ORF), 60 kDa (42-kDa ORF), and 42 kDa (22-kDa ORF). (**D**) Read coverage across a representative 60-kb window of the *E. coli* genome for the FZZ-tagged expression vectors, in the presence (+) and absence (-) of the corresponding untagged double-expression vectors (‘Dbl’). The y-axis represents counts per million (CPM), and the axis limit is set to 100.

**Figure S4.**
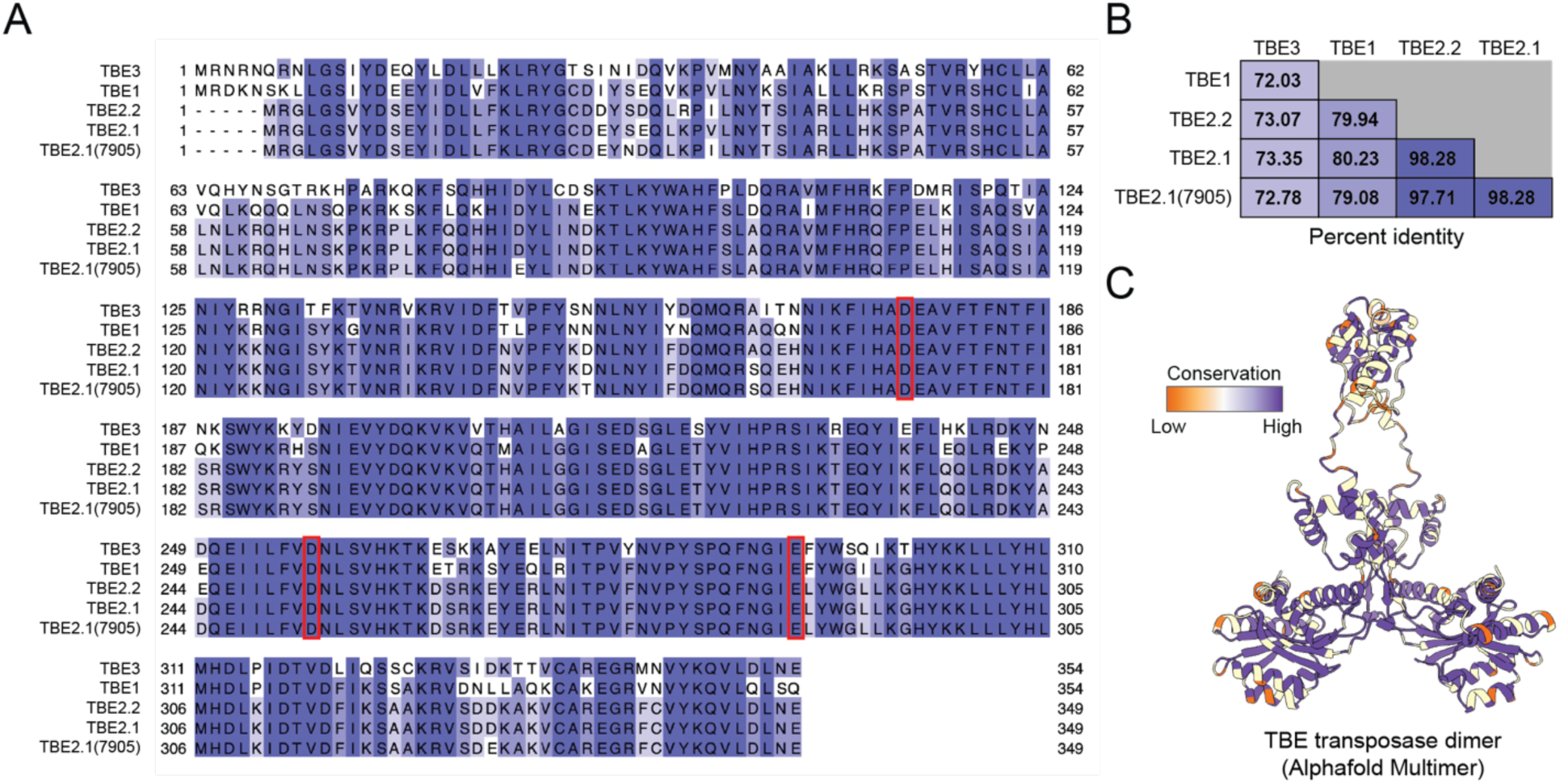
Alignment and amino acid similarity across TBE transposase families. (**A**) Multiple sequence alignment of the four TBE family transposase consensus sequences (Chen *et al*. 2016), alongside representative TBE2.1 family member (7905). Red boxes indicate residues belonging to the DDE catalytic triad. (**B**) Sequence identity matrix of the percent amino acid identity for the four TBE family consensus sequences and TBE2.1(7905). (**C**) TBE transposase dimer structural prediction colored by sequence conservation across the TBE families.

**Figure S5.**
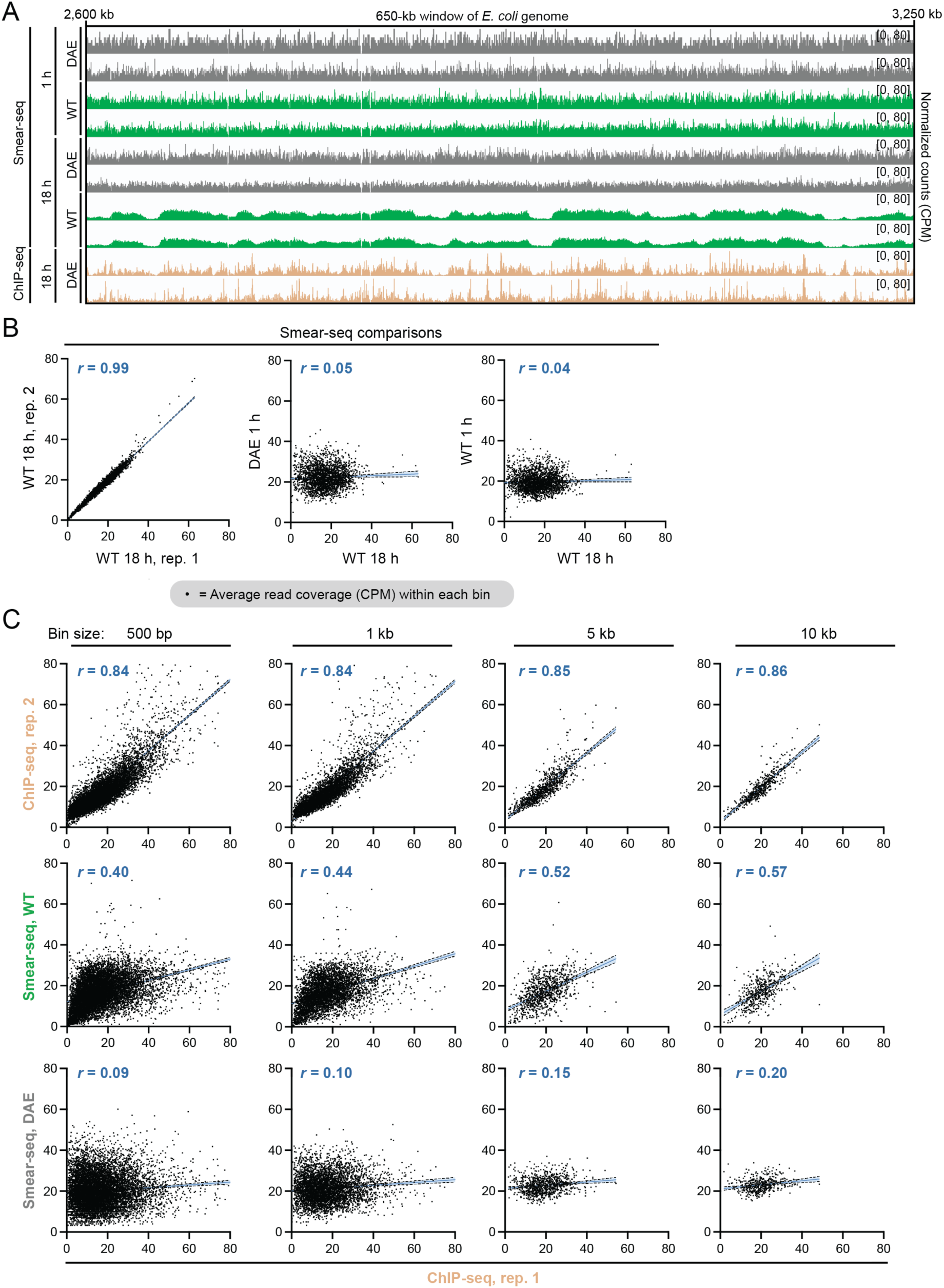
Comparison between ChIP-seq and Smear-seq datasets. (**A**) Read coverage profiles of all Smear-seq samples and the 18 h ChIP-seq DAE sample, in a 650-kb window of the *E. coli* genome. The y-axis represents CPM with the axis limit set to 80. (**B**) Comparisons between average read coverage in CPM across 2-kb bins of the genome. The Smear-seq signal at 18 h is strongly correlated across independent biological replicates. (**C**) Coverage comparisons when bin sizes are varied between 500 bp and 10 kb. *r* values represent the Pearson correlation, and linear regressions are shown with 95% confidence bands of the best-fit line.

**Figure S6.**
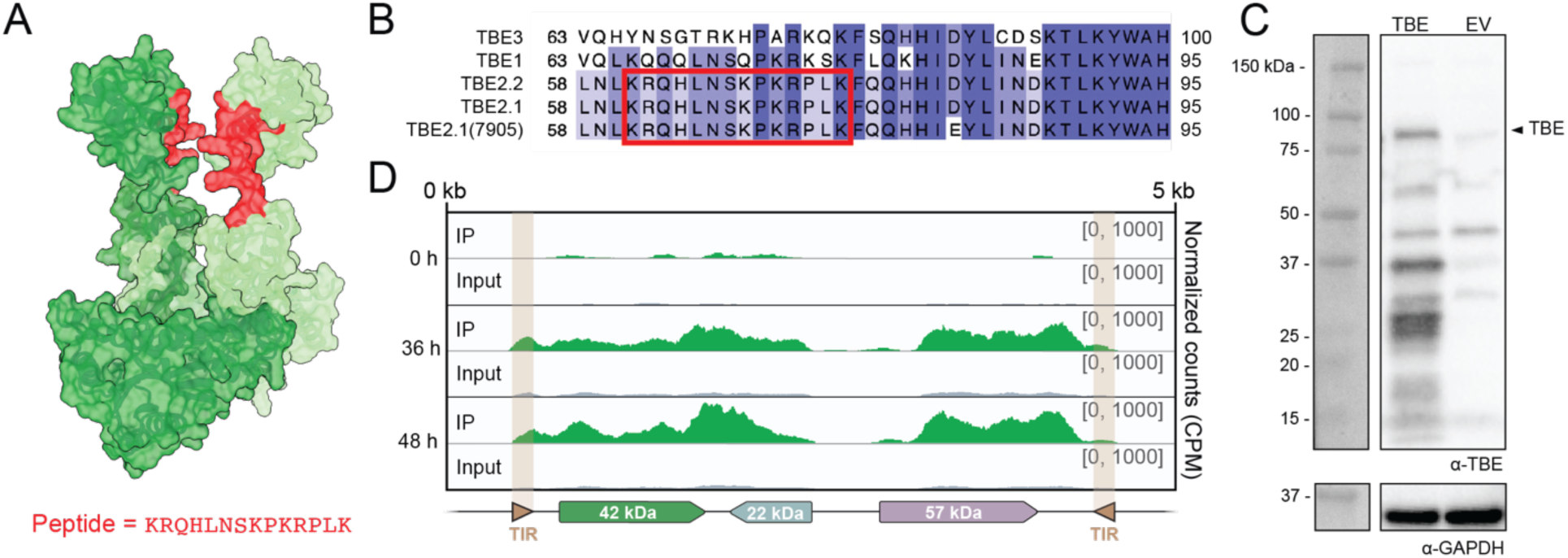
Generating and validating an anti-TBE transposase antibody. (**A**) Structural prediction of the TBE2.1 transposase dimer by AlphaFold-Multimer (Evans *et al*., 2022); red coloring indicates the predicted surface-exposed region that was selected for peptide synthesis and antibody generation. (**B**) Partial multiple sequence alignment of the four TBE-family transposase consensus sequences (from **Figure S4A**). Red box indicates the 14 residues selected for peptide synthesis. (**C**) Western hybridization to the anti-TBE transposase antibody (top) and anti-GAPDH antibody loading control (bottom). The two samples are *E. coli* BL12(DE3) cells expressing either the MBP-tagged 42-kDa plasmid (TBE) or an empty vector control (EV). The expected size of the MBP-tagged 42-kDa ORF is 88.6 kDa. (**D**) Read coverage mapped across the representative TBE2.1(7905) element demonstrates transposon localization, concentrated at genic regions, at 36 and 48 h. The y-axis represents counts per million (CPM), and the axis limit is set to 1000.

